# Cosine Similarity Estimation Using FracMinHash: Theoretical Analysis, Safety Conditions, and Implementation

**DOI:** 10.1101/2024.05.24.595805

**Authors:** Mahmudur Rahman Hera, David Koslicki

## Abstract

**Motivation:** The increasing number and volume of genomic and metagenomic data necessitates scalable and robust computational models for precise analysis. Sketching techniques utilizing *k*-mers from a biological sample have proven to be useful for large-scale analyses. In recent years, FracMinHash has emerged as a popular sketching technique and has been used in several useful applications. Recent studies on FracMinHash proved unbiased estimators for the containment and Jaccard indices. However, theoretical investigations for other metrics, such as the cosine similarity, are still lacking.

**Theoretical contributions:** In this paper, we present a theoretical framework for estimating cosine similarity from FracMinHash sketches. We establish conditions under which this estimation is sound, and recommend a minimum scale factor *s* for accurate results. Experimental evidence supports our theoretical findings.

**Practical contributions:** We also present `frac-kmc`, a fast and efficient FracMinHash sketch generator program. `frac-kmc` is the fastest known FracMinHash sketch generator, delivering accurate and precise results for cosine similarity estimation on real data. We show that by computing FracMinHash sketches using `frac-kmc`, we can estimate pairwise cosine similarity speedily and accurately on real data. `frac-kmc` is freely available here: https://github.com/KoslickiLab/frac-kmc/.

**ACM Subject Classification:** Applied computing → Computational biology

## 1 Introduction

With the growing number of reference genomes and the exponential increase in genomic and metagenomic data production, there is a critical need for the development of computational models that are both scalable and robust, as well as ensure precision in analysis. *k*-mer-based algorithms, particularly those utilizing sketching methods, are becoming increasingly popular for large-scale sequence analysis and metagenomic applications. A *k*-mer is a sequence of *k* consecutive nucleotides extracted from a longer sequence. Algorithms designed to work with *k*-mers decompose a long sequence into small *k*-mers and analyze based on the number of shared or dissimilar *k*-mers among multiple samples. Given the potentially vast number of distinct *k*-mers in a sequencing sample, sketching methods create a fingerprint of the *k*-mers (called a sketch) to work with these smaller sets, thereby reducing computational resource consumption. The most widely used sketching method for many years has been MinHash [3], originally introduced for document comparisons. `Mash` [17] was developed to apply MinHash to genomic data and has been extensively utilized. However, recent studies have indicated that MinHash sketches perform relatively poorly when comparing sets of very dissimilar sizes [13, 12]. Researchers have proposed various adjustments to MinHash to address this issue [2, 10, 12, 16]. One such example is the recently introduced FracMinHash sketch, which uses a variable sketch size instead of MinHash’s fixed-size scheme. FracMinHash was first introduced and used in the software sourmash [4, 19]. In simple words, a FracMinHash sketch retains *s* (0 ≤ *s* ≤ 1) fraction of the input set of *k*-mers. The scale factor *s* is a tunable parameter of the FracMinHash sketching technique, controlling the size of the generated sketch.

The first theoretical analysis of FracMinHash was introduced in [8], which showed how to obtain an unbiased estimator of the containment and the Jaccard indices computed using FracMinHash sketches. This work laid the theoretical foundation for calculating average nucleotide identity (ANI) via FracMinHash sketches and led to useful applications, such as ANI estimation in metagenomes [21], obtaining taxonomy off of metagenome samples [9], obtaining a functional profile from metagenomes [7], etc. Besides the Jaccard and the containment indices, there are other qualitative and quantitative metrics used in the literature when comparing two samples, such as cosine similarity, Bray-Curtis dissimilarity, KL divergence, Whittaker distance etc. Although these metrics are useful, and often used by researchers, a theoretical analysis of these metrics in the context of FracMinHash sketching is still missing – both in whether and when we can estimate the distance from FracMinHash sketches. In this paper, we present this theoretical analysis for the cosine similarity. We first explore the conditions when estimating cosine similarity from FracMinHash sketches is theoretically sound. We next show how these conditions can be used to recommend a minimum scale factor *s* that is theoretically safe to use when estimating cosine similarity from FracMinHash sketches. We supplement our theoretical findings with experimental evidence that show that these theoretical analyses are sound.

Apart from these theoretical results, our other contribution presented in this paper is implementing a fast and efficient FracMinHash sketch generator program, `frac-kmc`. Although FracMinHash sketches can readily be generated using the software `sourmash`, we found the program `sourmash` `sketch` to be slow for very large samples. Therefore, we developed `frac-kmc` by modifying a *k*-mer-counter tool `KMC` [5, 11, 6]. To the best of our knowledge, `frac-kmc` is the fastest FracMinHash sketch generator program when considering wall-clock time. We used `frac-kmc` to compute FracMinHash sketches and used the sketches to estimate cosine similarity values on real data, and found accurate and precise results. `frac-kmc` is freely available here: https://github.com/KoslickiLab/frac-kmc/. The analyses presented in this paper can be reproduced using the code here.

## 2 Preliminaries

We present the following preliminaries in their full generality, using generic notation such as Ω, a universal set. All theorems presented in Section 3 also hold for any universal set. In the case of sequence comparisons, the sets of interest, *A* and *B* are sets of *k*-mers, in the universe Ω = {*A, C, G, T* }^*k*^.

### FracMinHash sketching

Given a perfect hash function *h* : Ω → [0, *H*] for some *H* ∈ ℝ and a *scale factor s* where 0 ≤ *s* ≤ 1, a FracMinHash sketch of a set *A*, where *A* ⊆ Ω, is defined as follows:

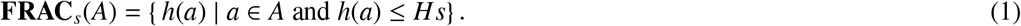

The scale factor *s* is a tuneable parameter that can modify the size of the sketch. For a fixed *s*, if the set *A* grows larger, the sketch **FRAC**_*s*_(*A*) grows proportionally.

### Vector form of a set

Let 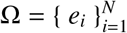 be a universal set, and let *A* ⊆ Ω. Then, the vector form of *A* is defined as follows: **u** = ⟨ *u*_*i*_ | *u*_*i*_ = 1 if *e*_*i*_ ∈ *A, u*_*i*_ = 0 if *e*_*i*_ ∉ *A* ⟩. In words, **u** is an *N*-dimensional vector, every entry representing the presence/absence of an element.

### Cosine similarity of two sets

Let *A* and *B* be two sets in Ω, and let the vector forms of *A* and *B* be **u** and **v**, respectively. Then, the cosine similarity of the sets *A* and *B* is defined as the cosine similarity of the vectors **u** and **v** as follows:

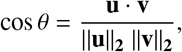

where **u** · **v** is the dot product of **u** and **v**. By using the triangle rule for cosine, we know the following:

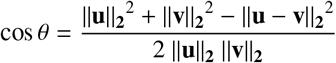

Throughout the paper, we have used the terms “cosine”, “cosine similarity”, and “similarity” analogously to mean the cosine similarity of two sets, unless stated otherwise.

### Chernoff bound for sum of Bernoulli random variables

Recall the classic Chernoff bounds: Let *X*_*i*_, *i* = 1, 2, …, *n* be *n* independent Bernoulli random variables.

If 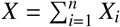 and *E*[*X*] = *µ*, then the following holds [15]

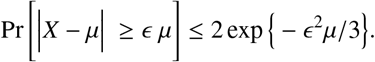

## 3 Theoretical Results

In this section, we present our theoretical findings. Ideally, if the cosine similarity of two sets *A* and *B* is cos θ, and the cosine similarity of **FRAC**_*s*_(*A*) and **FRAC**_*s*_(*B*) is cos θ^′^, then we want the following to hold in all cases with high probability:

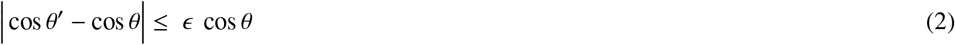

where *ϵ* → 0. In other words, the cosine similarity between two FracMinHash sketches approximates the cosine similarity between the original sets. In this ideal case, we would be able to compare two biological sequences and estimate the cosine similarity of the set of *k*-mers (extracted from the two sequences) by computing the cosine similarity of the FracMinHash sketches. Unfortunately, this does not hold in all cases. In this section, we present theoretical conditions where Equation (2) holds (and where it breaks down). For the sake of continuity, all proofs of the theorems are included in Section 5.

### ▸ Theorem 1.

*Let* 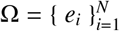 *be a given set (universe), and let A* ⊆ Ω. *If* **u** = ⟨ *u*_*i*_ | *u*_*i*_ = 1 *if e*_*i*_ ∈ *A, u*_*i*_ = 0 *if e*_*i*_ ∉ *A* ⟩ *and if* 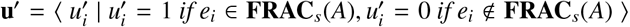, *then the following holds:*

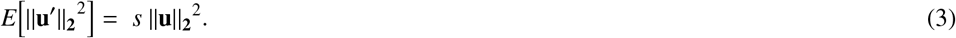

**Proof**. See Section 5.3. ◂

Theorem 1 quantifies the expected squared length of **u**^′^. We next show that the square of the length of **u**^′^ is well concentrated around this expected value.

### ▸ Theorem 2.

*Let* 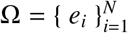 *be a given set (universe), and let A* ⊆ Ω. *If* **u** = ⟨ *u*_*i*_ | *u*_*i*_ = 1 *if e*_*i*_ ∈ *A, u*_*i*_ = 0 *if e*_*i*_ *∉ A* ⟩ *and if*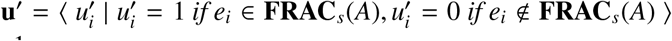, *then the following holds for* 0 < ϵ < 1:

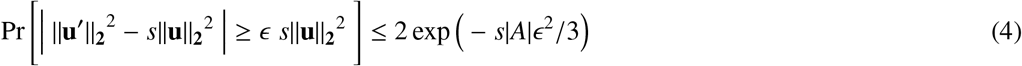

**Proof**. See Section 5.3.

Using this strong concentration of ||**u**^′^||_**2**_^2^ and ||**v**^′^||_**2**_^2^ around their respective mean values, we can quantify how well cos *θ*^′^ estimates the true cosine cos *θ*.

### ▸ Theorem 3.

*Let* 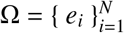 *be a given set (universe), and let A, B* ⊆ Ω *be two sets in the universe. Let the cosine similarity of the sets A and B be* cos *θ, and let* cos *θ*^′^ *be the cosine similarity of the sketched sets* **FRAC**_*s*_(*A*) *and* **FRAC**_*s*_(*B*). *Then, the following holds:*

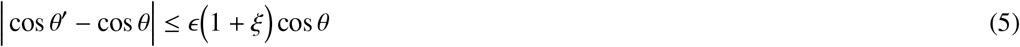

*with a probability of at least* 1−6 exp −*s* min(*m, n*) *ϵ*^2^/ 3, *where* 0 < *ϵ* < 1. *Here* |*A*| = *m*, |*B*| = *n*, |*A* ∩ *B*| = *q, and* ξ = 3(*m* + *n* − 2*q*)/*q*.

**Proof**. See Section 5.3. ◂

Theorem 3 indicates that the cosine similarity between two sketched sets approximates the cosine similarity of the original sets with approximation error being bounded by the relative similarity of the original sets. The other conditions describe when the error is properly bounded.

Before presenting the experimental results, we briefly discuss what *q* and ξ signify in Theorem 3. We first note that when *q* < (*m* + *n*)/(2 + *c*/3) for some positive constant *c*, then the right-hand side of Equation (5) becomes unbounded because ξ > *c*. In such a case, there is no theoretical guarantee that the error in estimating cosine using FracMinHash to be small. Indeed, as *q* → 0, we note that cos *θ* → 0, and intuitively, FracMinHash sketches **FRAC**_*s*_(*A*) and **FRAC**_*s*_(*B*) cannot likely retain the few common items in *A* and *B*. In such a case, Equation (2) does not hold for small ϵ. We do note that in this case, **FRAC**_*s*_(*A*) ∩ **FRAC**_*s*_(*B*) will have nearly zero elements, and as a result, cos *θ*^′^ → 0. The only limitation of Theorem 3 is that it does not guarantee that two near-zero quantities will be proportionally close. Nevertheless, estimating cosine using FracMinHash sketches will still work, since the cosine similarity of the sketched sets will also be near-zero.

On the other hand, ξ can be bounded by a positive constant *c* when *q* ≥ (*m* + *n*)/(2 + *c*/3), i.e. when there are enough common elements between *A* and *B*. In practical scenarios, this is the case as we compare a pair of biological sequences, and Theorem 3 gives guarantees that estimating cosine similarity using FracMinHash sketches is theoretically sound.

### 3.1 Suggesting a minimum scale factor *s*

We conclude our theoretical results section by suggesting a minimum scale factor that is theoretically safe to use. The probability guarantee in Theorem 3 allows us to recommend a scale factor *s* for a desired error rate *δ* and a desired level of confidence α, 0 ≤ α < 1. If we want to have a guarantee of at least α, 0 ≤ α < 1, that the estimated cosine cos *θ*^′^ will be in a 1 ± *δ* factor of the true cosine cos *θ* where 0 ≤ *δ* < 1, then we require a scale factor *s*, such that

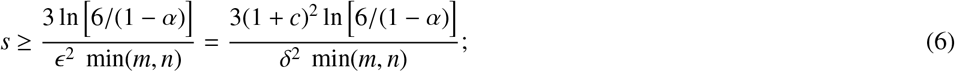

where *ξ* in Theorem 3 is bounded by a constant *c, c* > 0. If we want a higher level of confidence, or if we want a smaller window of error, we require a larger scale factor. The given sets *A* and *B* dictate the other terms – if there is a large number of elements in *A* ∩ *B*, then ξ is bounded by a smaller *c*, and a smaller scale factor *s* suffices. And finally, the larger the sets *A* and *B* are, the smaller scale factor *s* can be.

## 4 Experimental Results

In this section, we present our experimental results. We first show results supporting the theory we presented in Section 3. Then, we discuss a fast and efficient program to compute FracMinHash sketches from nucleotide sequences. We named this program `frac-kmc`. Finally, we present the performance of `frac-kmc` on real biological sequences.

### 4.1 Our suggested scale factors are safer to estimate cosine

We start by presenting what the scale factors suggested by Equation (6) look like for various α and *δ*. Table 1 shows various suggested scale factors when min(*m, n*) = 10K, and Table 2 shows suggested scale factors when min(*m, n*) = 10M. We notice that the theory accounts for a larger number of elements in the sets that are being compared against each other. With only 10K elements, if we want the estimated cosine to be within ± 5% of the original cosine, then the theory suggests that we have to use a scale factor of 1. In other words, there is no scope for sub-sampling at this desired resolution. It is only at *δ* ≥ 0.06 that we can get away with sub-sampling, although the recommended scale factor is not very small to be drastically helpful in reducing computational resources.

**Table 1.** Suggested scale factors for various levels of desired confidence and various tolerable rates of error, when min(*m, n*) = 10000. For only 10K elements, if the tolerable error is up to 5%, we cannot but use all elements to get the desired accuracy.

| Tolerable Error, $\delta$ | Desired level of confidence, $\alpha$ | | | | |
| --- | --- | --- | --- | --- | --- |
|  | 0.91 | 0.93 | 0.95 | 0.97 | 0.99 |
| 0.01 | 1.0000 | 1.0000 | 1.0000 | 1.0000 | 1.0000 |
| 0.03 | 1.0000 | 1.0000 | 1.0000 | 1.0000 | 1.0000 |
| 0.05 | 1.0000 | 1.0000 | 1.0000 | 1.0000 | 1.0000 |
| 0.07 | 0.5785 | 0.6132 | 0.6595 | 0.7299 | 0.8812 |
| 0.09 | 0.3500 | 0.3709 | 0.3990 | 0.4415 | 0.5331 |
| 0.1 | 0.2835 | 0.2835 | 0.3232 | 0.3576 | 0.4318 |

**Table 2.** Suggested scale factors for various levels of desired confidence and various tolerable rates of error, when min(*m, n*) = 10000000. For 10M elements, we can use a small fraction of the elements to get the desired accuracy when estimating the cosine similarity.

| Tolerable Error, $\delta$ | Desired level of confidence, $\alpha$ | | | | |
| --- | --- | --- | --- | --- | --- |
|  | 0.91 | 0.93 | 0.95 | 0.97 | 0.99 |
| 0.01 | 0.0283 | 0.0300 | 0.0323 | 0.0358 | 0.0432 |
| 0.03 | 0.0031 | 0.0033 | 0.0036 | 0.0040 | 0.0048 |
| 0.05 | 0.0011 | 0.0012 | 0.0013 | 0.0014 | 0.0017 |
| 0.07 | 0.0006 | 0.0006 | 0.0007 | 0.0007 | 0.0009 |
| 0.09 | 0.0003 | 0.0004 | 0.0004 | 0.0004 | 0.0005 |
| 0.1 | 0.0003 | 0.0003 | 0.0003 | 0.0004 | 0.0004 |

On the other hand, as documented in Table 2, we can use a scale factor of roughly 0.0003 ∼ 0.0004 to allow a 10% window for error when we have at least 10M elements. If we want to be *very* accurate and only allow 1% error, we need to obtain FracMinHash sketches with a scale factor of roughly 0.03 ∼ 0.04. From Table 2, we also notice that with a higher level of desired confidence *α*, we need to employ larger scale factors; although the effect a larger α has on the suggested scale factor is less prominent than the effect of a smaller *δ*.

In all these cases, we used *c* = 0.5, which was arbitrarily assumed that ξ is bounded by 0.5. We experimented with other values of *c* and noticed similar patterns. On simulated data, the scale factors suggested for *c* = 0.5 give satisfactory results, as presented below.

We next show the usefulness of using these recommended scale factors, in contrast to a preset value. The state-of-the-art program to compute and analyze FracMinHash sketches is sourmash [4, 19], which uses a default scale factor of 1/1000. As a result, many studies that use sourmash use this default value, even though the tool can work with other non-default scale factors. We show that a preset scale factor may result in an error higher than expected. In this set of experiments, we simulated a universe of 1M elements. We then randomly selected two sets *A* and *B* from this universe. We varied the number of elements in these sets from 100K to 500K. The actual elements were selected randomly. We then calculated the true cosine of *A* and *B* using all elements. After that, we used the preset scale factor of 1/1000 to compute FracMinHash sketches of *A* and *B*, and estimated the cosine using these sketches. We also set the tolerable error rate (*δ*) at 5%, and desired level of confidence (*α*) at 95%. We then computed FracMinHash sketches of *A* and *B* using the scale factor suggested by Equation (6), and used these sketches to estimate the cosine. We then recorded if these estimated cosine values fall in the range cos *θ* (1 ± 0.05). We repeated the experiment 1000 times for all sizes of *A* and *B*. Table 3 shows the fraction of times the similarities estimated using a fixed scale factor of 1/1000 fall within ±5% of the true cosine. Table 5 shows the fraction of times the similarities estimated using the scale factor recommended by Equation (6) fall within ±5% of the true cosine. The list of suggested scale factors in these scenarios is shown in Table 4.

**Table 3.** Fraction of times the estimated cosine falls within ±5% of the true cosine of *A* and *B*, for different sizes of *A* and *B*. The similarities were estimated **using a scale factor of 1***/***1000**, which is the default in `sourmash`. In a large fraction of times, the estimated cosine is *not* within ±5% of the true cosine.

| Num. elements in B | Num. elements in A |  |  |  |  |
| --- | --- | --- | --- | --- | --- |
|  | 100K | 200K | 300K | 400K | 500K |
| 100K | 0.09 | 0.20 | 0.25 | 0.32 | 0.38 |
| 200K | 0.17 | 0.29 | 0.55 | 0.47 | 0.51 |
| 300K | 0.28 | 0.40 | 0.54 | 0.58 | 0.57 |
| 400K | 0.32 | 0.45 | 0.61 | 0.71 | 0.74 |
| 500K | 0.45 | 0.42 | 0.73 | 0.66 | 0.83 |

**Table 4.**
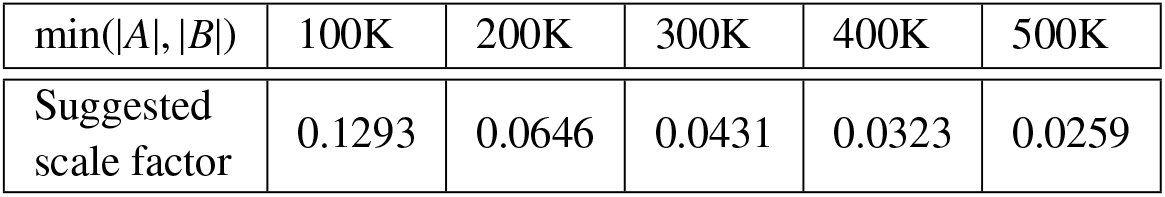
Suggested scale factors for various min(|*A*|, |*B*|), as calculated by Equation (6). α = 0.95, *δ* = 0.05, and *c* = 0.5 was used.

**Table 5.**
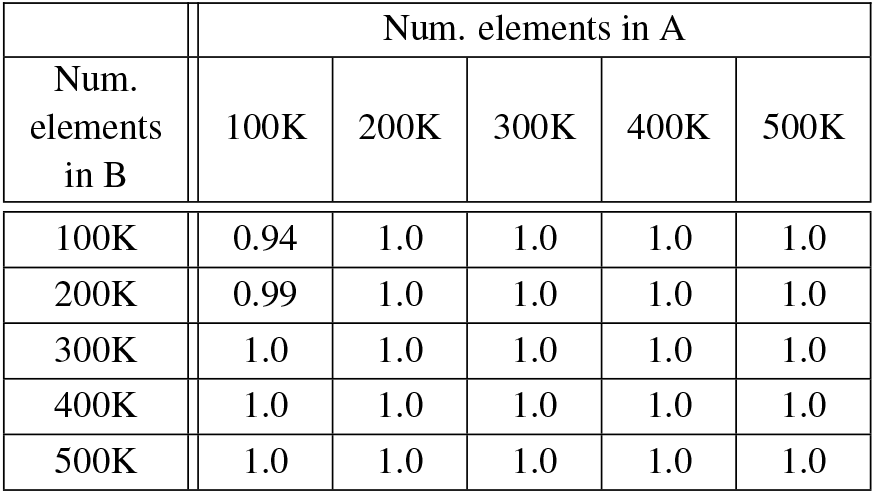
Fraction of times the estimated cosine falls within ±5% of the true cosine of *A* and *B*, for different sizes of *A* and *B*. The similarities were estimated **using the scale factor suggested by Equation (6)**. In almost all instances, the recommended scale factor can estimate the similarity so that the estimated value is within ±5% of the true similarity.

| Num.<br>elements<br>in B | Num. elements in A |  |  |  |  |
| --- | --- | --- | --- | --- | --- |
|  | 100K | 200K | 300K | 400K | 500K |
| 100K | 0.94 | 1.0 | 1.0 | 1.0 | 1.0 |
| 200K | 0.99 | 1.0 | 1.0 | 1.0 | 1.0 |
| 300K | 1.0 | 1.0 | 1.0 | 1.0 | 1.0 |
| 400K | 1.0 | 1.0 | 1.0 | 1.0 | 1.0 |
| 500K | 1.0 | 1.0 | 1.0 | 1.0 | 1.0 |

These results clearly show the usefulness of the recommended scale factors over a preset value. Almost 100% of times, the recommended scale factor can estimate a cosine within the tolerable error range, whereas using a preset scale factor can result in a larger error. We computed the recommended scale factor using *c* = 0.5. This is equivalent to |*A* ∩ *B*| ≥ ∼ 46% of |*A*| + |*B*|. Despite this being a non-realistic assumption for distantly related samples, the suggested scale factors still perform well because of the many pessimistic steps in Theorem 3, which states conditions that are more stringent than necessary. Nonetheless, it is clear that using the default scale factor of 1/1000 may not be well-suited where a higher resolution around the true value is required.

### 4.2 frac-kmc computes FracMinHash sketches faster

After establishing the conditions when FracMinHash sketches can be safely used to estimate the cosine similarity, we next wanted to use FracMinHash sketches on real biological sequences. Ideally, we wanted to show that by using FracMinHash sketches, we can compute the pairwise similarity matrix for a number of sequences faster than tools that use all *k*-mers. The fastest tool that can compute pairwise similarity/distance matrix from a list of sequences is currently `Simka` [1], whereas the state-of-the-art tool to compute FracMinHash sketches is `sourmash` [4, 19]. Naturally, we tried to use sourmash to first compute FracMinHash sketches, and later compare the sketches to obtain a pairwise similarity matrix. Unfortunately, we found that the command that computes FracMinHash sketches (called `sourmash sketch`) is many times slower than `Simka`. We noted that this is because `sourmash` treats input sequence files in a serialized manner, where there is scope for parallelism over multiple threads to make the processing faster. This is surprising since *k*-mer-counting is a very well-studied problem in the literature.

Therefore, for practical purposes, we decided to write a new FracMinHash sketch generator program by modifying the source code of a fast and efficient *k*-mer-counter `KMC` [11]. Details of `frac-kmc` are included in Section 5.1. Figure 1 shows a running time comparison for the commands `sourmash sketch` and `frac-kmc sketch` on files of different sizes. The files are compressed `fastq.gz` files, which were randomly selected from the Human Microbiome Project [18]. We verified that sketches produced by the two programs are identical by running `sourmash compare`. The comparison shows that `frac-kmc` consistently runs ∼10 times faster than `sourmash` to sketch input files. For this set of analyses, we used the latest version of sourmash: 4.8.8, as of 1 May 2024. We used 128 threads to run `frac-kmc`. Both `sourmash` and `frac-kmc` were run to not keep track of abundances. We ran the programs to compute sketches for *k* = 21 and scale factor *s* = 1/1000. We tested with other values of *k* and *s* and saw similar results.

**Figure 1.**
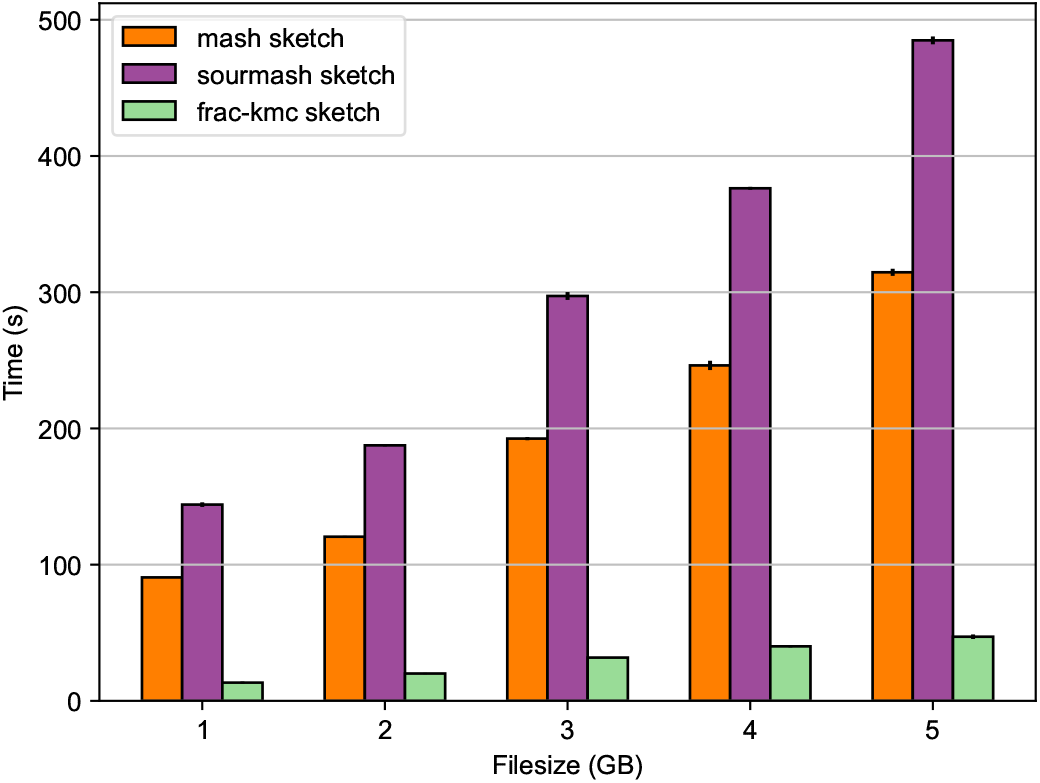
Wall-clock time required by the commands `mash sketch`, `sourmash sketch` and `frac-kmc-sketch` to compute a sketch. The input files are `fastq.gz` files containing metagenome samples taken from the human gut. MinHash sketches of 1000 was computed, and FracMinHash sketch with scale factors *s* = 0.001 was computed. `frac-kmc` finishes ∼ 10 times faster than `sourmash`, and ∼ 6.7 times faster than `Mash`.

Figure 1 also shows the average running time to compute MinHash sketches (sketch size = 1000 hashes) from the same files using `Mash` (version 2.0). Both `Mash` and `sourmash` run slower than `frac-kmc` mainly because of processing input files in a serialized manner. When working on many input files, both `Mash` and `sourmash` can parallelize the workload by processing one file on every core. Nevertheless, using `frac-kmc` can still be helpful to end users, especially for processing very large files, when the difference in running time is starkly observed.

### 4.3 frac-kmc estimates cosine similarity accurately

We next show that by using FracMinHash sketches computed by `frac-kmc`, we can estimate cosine similarity faster than `Simka` [1] (which uses all *k*-mers to operate), and more accurately than `Mash` [17] (which uses fixed size MinHash sketches). For this set of experiments, we used two datasets: the **Ecoli** dataset contains 3682 E.coli genome assemblies, and the **HMP** dataset contains 300 metagenome samples collected from the human gut, taken from the Human Microbiome Project [18]. We ran `Simka`, `Mash`, and `frac-kmc` on these datasets to produce the pairwise cosine similarity matrix. Details of the datasets, and how the programs were run are included in Section 5.2.

Total wall-clock time and CPU time to compute pairwise cosine for all three tools are shown in Figure 2. For both the **Ecoli** and the **HMP** dataset, we randomly selected a number of samples and ran the tools. We found that as the number of samples gets to roughly 125, `Simka` does not exit even after letting it run for more than 48 hours. Other than these extremes, the running time of `Simka` grows linearly with the number of samples (top-left and mid-left plots of Figure 2), and the tool naturally requires more resources to keep track of all *k*-mers. We found that `Simka` operates by creating many `SimkaCount` and `SimkaMerge` processes, which are not spawned as descendants of the mother `Simka` process. Therefore, we found no good way to measure the CPU time consumed by `Simka`, and are not including the CPU time comparisons for the smaller number of samples.

**Figure 2.**
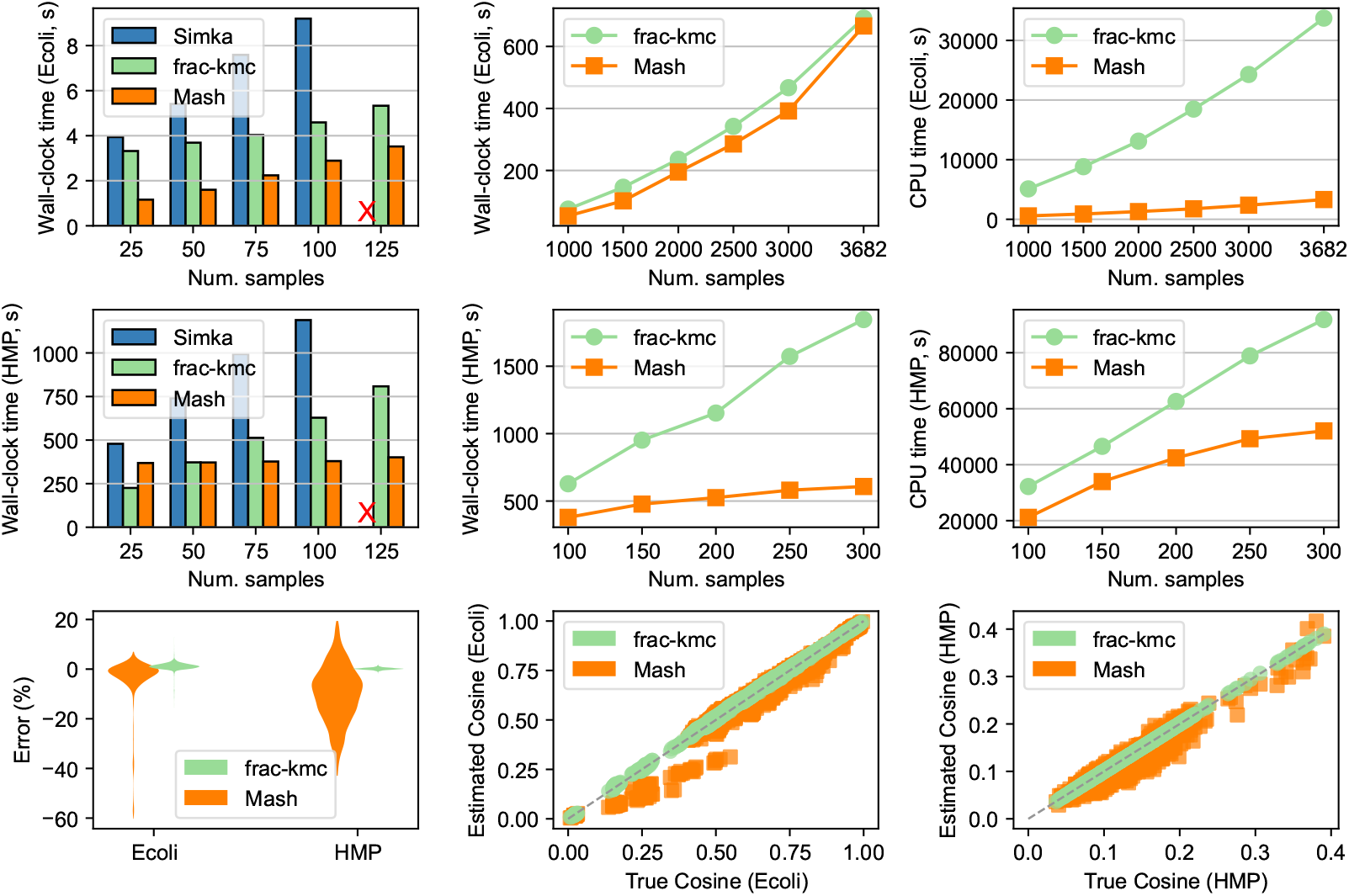
Running time and accuracy of the tools on **Ecoli** and **HMP** datasets. Top row shows total wall-clock time and CPU time to run the tools on the **Ecoli** dataset. The top-left plot shows wall-clock time for all three tools when running on 25-125 randomly selected samples from the dataset. An X indicates that Simka did not exit after > 48 hours. The top-center and the top-right plots show total wall-clock time and total CPU time, respectively, for `frac-kmc` and Mash on the **Ecoli** dataset for a larger number of samples (to which, Simka does not scale). The middle row shows the same plots for the **HMP** dataset. The bottom row shows the accuracy of FracMinHash and MinHash sketches in estimating cosine similarities. The bottom-left plot shows the distributions of error percentages. The bottom-center and bottom-right plots show estimated vs. true cosine values for the **Ecoli** (3682 samples) and the **HMP** (100 samples) dataset, respectively.

To explain the running time of `Mash` and `frac-kmc`, we note that there are two main stages to running these tools: computing sketches from input files, and using sketches to estimate cosine values. As presented in Figure 1, `frac-kmc` is faster than `Mash` in wall-clock time for the first stage: computing sketches, although `frac-kmc` requires more CPU time since it runs on multiple cores. The code that executes the second stage is identical for the two tools. The time required in this stage is therefore determined by the sketch sizes. Therefore, in this stage, using `frac-kmc` requires more wall-clock time as well as CPU time. Indeed, as we push the number of samples to higher values, we observe that using `Mash` uses less CPU time, as shown in the top-right and mid-right plots in Figure 2 (roughly 8.9-10.1x less CPU time than frac-kmc in the **Ecoli** dataset, and roughly 1.37x-1.76x less CPU time in the **HMP** dataset).

Next, we turn to explain the wall-clock times in running `Mash` and `frac-kmc`. Note that the input metagenome files in the **HMP** dataset are much larger than the genome files in the **Ecoli** dataset. Therefore, the FracMinHash sketches computed for the **HMP** samples are also much larger. Consequently, in the **HMP** dataset, the calculation of pairwise metrics dominates the overall running time, and hence `frac-kmc` takes longer to run in the **HMP** dataset, as evident in the mid-center plot of Figure 2. On the other hand, the FracMinHash sketches are closer to the MinHash sketches in size for the **Ecoli** dataset. Therefore, the wall-clock time of `frac-kmc` in the **Ecoli** dataset is much closer to `Mash`, as seen in the top-middle plot of Figure 2.

We also present the accuracy of `Mash` and `frac-kmc` outputs with respect to the true cosine values (calculated using all *k*-mers), in the bottom row of Figure 2. Here, the results are shown for all 3682 samples in the **Ecoli** dataset, and only 100 samples in the **HMP** dataset. We calculated the true cosine values for all pairs in the **Ecoli** dataset by manually counting all the *k*-mers. For the **HMP** dataset, doing so was not feasible, and therefore, we did this analysis for only 100 samples (the largest number of samples to which `Simka` scales). In the bottom-left plot of Figure 2, we show the distribution of percentage of errors when we use `frac-kmc` and `Mash` to estimate pairwise cosine values. The plot shows that while the percentage of error for both `Mash` and `frac-kmc` are close to zero in expectation for the **Ecoli** dataset, we get a higher variability in error by using `Mash`. In the **HMP** dataset, the samples are just too large for `Mash` to even give an expected error of 0. On the other hand, with larger sketches, FracMinHash gives an almost perfect estimation. The bottom-center and bottom-right plots show estimated vs. true cosine values for two datasets. The bottom-center plot shows that using `frac-kmc` gives a better estimate across the entire range of 0 to 1. Although the **HMP** dataset does not have true cosine values covering the entire range, using `frac-kmc` gives a better estimate, as evident in the bottom-right plot.

## 5 Methods

### 5.1 Implementation of frac-kmc

The core motivation behind implementing `frac-kmc` was that sourmash sketch dna was very slow for larger files. Therefore, we decided to use a fast and efficient *k*-mer-counting program. There are many *k*-mer-counters available in the literature, namely `jellyfish` [14], `DSK` [20], `KMC` [11] etc. We decided to use `KMC` since its source code was easy to understand and navigate. Instead of running `KMC` and iterating through all *k*-mers in `KMC`’s output, we decided to modify the source code so that only the *k*-mers in the sketch were retained in the output. This made the entire program many times faster since typical scale factors used to compute FracMinHash sketches are very small. Therefore, we implemented the 64-bit MurMurHash function in C++ within the source code of `KMC`, and made the necessary changes so that instead of keeping track of all the *k*-mers, the program now kept track of only the *k*-mers whose hash value fell below the cut-off threshold. As a result, the succinct *k*-mer-database constructed by this modified `KMC` now contained only the relevant *k*-mers.

Finally, we modified the program `kmc dump` so that instead of writing all the *k*-mers in an output file, it now wrote the 64-bit MurMurHash values for the kmers in a sorted list – which is the output format of `sourmash sketch`. We named this program `frac-kmc`. After generating sketches from the same file using `frac-kmc` and `sourmash`, we used `sourmash compare` to confirm that the sketches are identical.

### 5.2 Generating results in Figure 2

#### 5.2.1 Datasets

The datasets we used are:

1. **Ecoli:** We collected all 3682 E. coli genome assemblies in NCBI.
2. **HMP:** We collected whole genome shotgun sequences from the Human Microbiome Project [18]. We randomly selected 300 gzipped fastq files corresponding to samples collected from the human gut.

The metagenome samples in the **HMP** dataset have an average file size of 1.88 GB and a median file size of 1.72 GB. The smallest file size is 58 MB, and the largest file size is 5.5 GB. This dataset works as a stress test for all the tools, where the input files are very large, reflecting real-life metagenome data; although the number of total samples is manageable. On the other hand, the **Ecoli** dataset challenges all the tools because the number of samples is very large (there are roughly 67 million pairs), although every individual file is quite small and easy to process.

#### 5.2.2 Running Simka, Mash, and frac-kmc

We ran Simka, `Mash`, and `frac-kmc` on the **Ecoli** and the **HMP** dataset, to produce the pairwise cosine similarity matrix. Simka readily produces several similarity and dissimilarity metrics when invoked on a list of input files. However, it does not produce cosine similarity. Therefore, we took the Chord distances generated by `Simka` and converted them to cosine similarities.

We used `Mash` and `frac-kmc` to compute MinHash and FracMinHash sketches of the input files, respectively. We then used a parallelized program to read all the sketches and compute the cosine similarity using the sketches. We ran `Mash` to generate MinHash sketches of size 1000, the default value. We also experimented with larger MinHash sketch sizes but found that although using larger sketches consumes more resources, it does not improve accuracy. The minimum number of *k*-mers in all files we used was roughly 4.8 million. In such a case, the minimum scale factor suggested by Equation (6) is 0.0006 (using *c* = 0.5, *δ* = 10%, *α* = 0.95). Therefore, we simply used the `sourmash` default value, 1/1000 to generate the FracMinHash sketches when running `frac-kmc`. All three tools were run on 128 cores of the same machine, including the multi-threaded code segment that reads MinHash and FracMinHash sketches, computes pairwise cosine similarity values, and writes them into an output file.

### 5.3 Proofs of theorems

#### ▸ Theorem 1.

*Let* 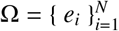 *be a given set (universe), and let A* ⊆ Ω. *If* **u** = ⟨ *u*_*i*_ | *u*_*i*_ = 1 *if e*_*i*_ ∈ *A, u*_*i*_ = 0 *if e*_*i*_ <∉ *A* ⟩ *and if*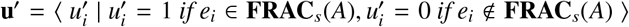, *then the following holds:*

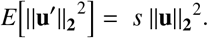

**Proof**. Let *I*_*i*_ be an indicator variable as follows:

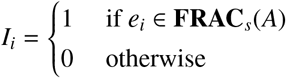

for all *i* such that *e*_*i*_ ∈ *A*. We note that *E*[*I*_*i*_] = Pr[*I*_*i*_ = 1] = *s*. We also observe that if *I*_*i*_ = 1, then 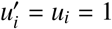.

Using these facts, we have the following:

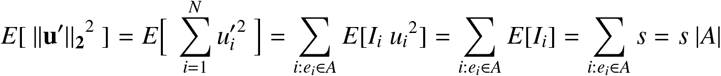

Of course, **u** being a binary vector, ||**u**||_**2**_^2^ = |*A*|, which completes the proof.

#### ▸ Theorem 2.

*Let* 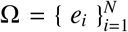*be a given set (universe), and let A* ⊆ Ω. *If* **u** = ⟨ *u*_*i*_ | *u*_*i*_ = 1 *if e*_*i*_ ∈ *A, u*_*i*_ = 0 *if e*_*i*_ *∉ A* ⟩ *and if* 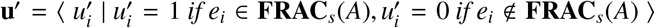⟩, *then the following holds for* 0 < *ϵ* < 1:

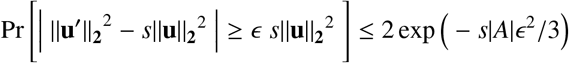

**Proof**. Using the same indicator *I*_*i*_ used in the proof of Theorem 1, we note the following:

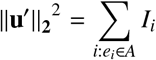

Therefore, ||**u**^′^||_**2**_^2^ is simply a sum of Bernoulli random variables. This allows for the use of the Chernoff concentration inequality for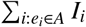. Using the fact that |*A*| = ||**u**||_**2**_^2^ completes the proof.

#### ▸ Theorem 3.

*Let* 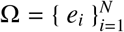 *be a given set (universe), and let A, B* ⊆ Ω *be two sets in the universe. Let the cosine similarity of the sets A and B be* cos *θ, and that of the sets* **FRAC**_*s*_(*A*) *and* **FRAC**_*s*_(*B*) *be* cos *θ*^′^. *Then, the following holds:*

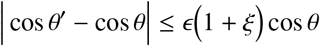

*with a probability of at least* 1−6 exp {−*s* min(*m, n*) ϵ^2^/ 3}, *where* 0 < ϵ < 1. *Here* |*A*| = *m*, |*B*| = *n*, |*A* ∩ *B*| = *q, and* ξ = 3(*m* + *n* − 2*q*)/*q*.

**Proof**. Let us define **u** and **u**^′^ as follows: **u** = ⟨ *u*_*i*_ | *u*_*i*_ = 1 if *e*_*i*_ ∈ *A, u*_*i*_ = 0 if *e*_*i*_ *∉ A* ⟩, and 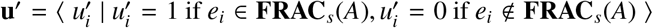. Let us also define **v** and **v**^′^ for the sets *B* and **FRAC**_*s*_(*B*) in an analogous manner, respectively.

We prove Theorem 3 by proving the following claims.

▹ Claim 4. With high probability, the following holds for 0 < ϵ < 1:

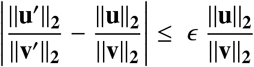

Proof. From Theorem 2, we know that the following holds:

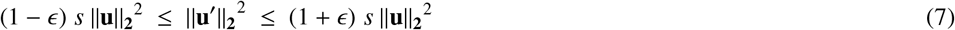

with a probability of at least 1 − 2 exp{−*s m ϵ*^2^/3}, for 0 < *ϵ* < 1.

Similarly, we also know the following holds:

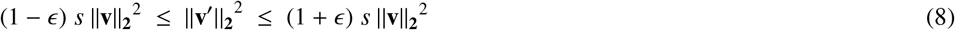

with a probability of at least 1 − 2 exp{−*s n ϵ*^2^/3}, for 0 < *ϵ* < 1.

Using these facts, and by taking square root and ratio, the following holds:

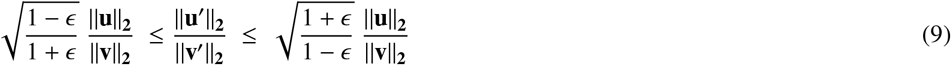

with a probability of at least 1 − 4 exp{−*s* min(*m, n*) *ϵ*^2^/3}, where 0 < *ϵ* < 1. The probability was calculated using a union bound of the probabilities of Equation (7) and Equation (8). We conclude the proof by using the fact that 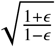 is simply 1 + *ϵ* + **O**(ϵ^2^), where for small *ϵ*, **O**(*ϵ*^2^) is dominated by *ϵ*. A similar argument for 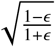 allows us to write the following:

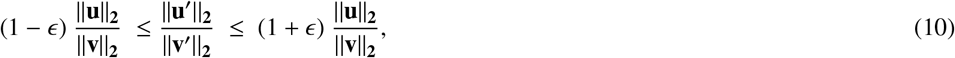

which completes the proof.

▹ Claim 5. With high probability, the following is true for 0 < *ϵ* < 1.

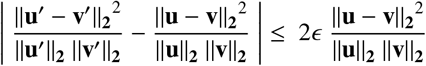

Proof. From Theorem 2, by plugging in **u** − **v** at the place of **u**, we know the following holds with high probability, where 0 < *ϵ* < 1.

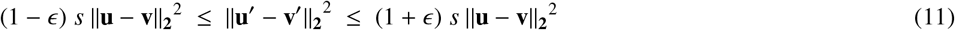

We next divide Equation (11) by the square root of Equation (7) and Equation (8). This gives us the following for 0 < *ϵ* < 1:

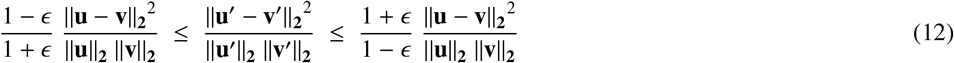

By union bound, this holds with a probability of at least 1 − 6 exp{−*s* min(*m, n*) *ϵ*^2^/3}, where 0 < *ϵ* < 1.

Finally, we note that 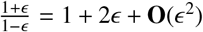 when 0 < *ϵ* < 1. For small *ϵ*, **O**(*ϵ*^2^) is dominated by 2*ϵ*. A similar argument for 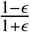 concludes the proof.

With the facts established in Claim 4 and Claim 5, we now calculate the following:

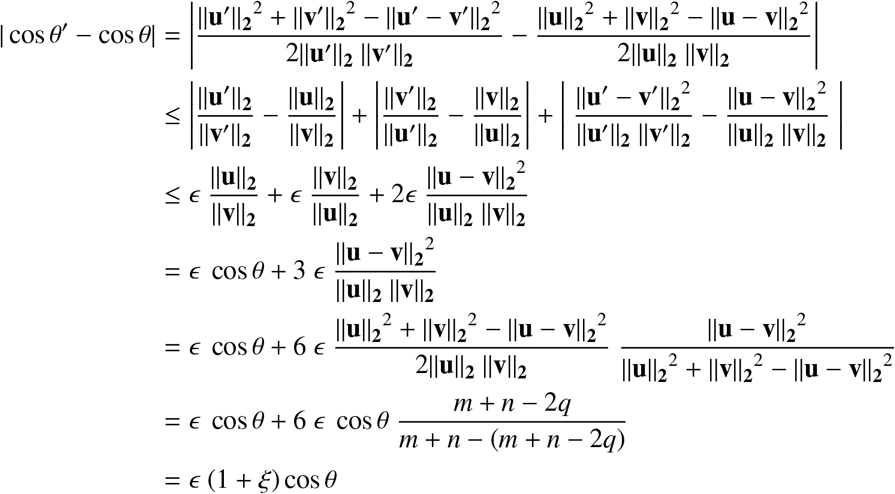

where *ξ* = 3(*m* + *n* − 2*q*)/*q*. In the derivation above, we have used the cosine triangle rule (elaborated in Section 2) in the first step, |*a* + *b*| ≤ |*a*| + |*b*| in the second step, and the cardinality of the sets *A, B*, and (*A* ∪ *B*) \ (*A* ∩ *B*) in the second to last step. Finally, the probability with which the above holds is the same as Claim 5 – which concludes the proof.

## 6 Discussions

### 6.1 Conclusions

Sketching-based methods allow practitioners to lower computational resource usages many-fold while keeping the accuracy reasonably well. In this paper, we analyzed such a sketching technique, FracMinHash, in estimating the cosine similarity via the sketches. We analyzed the conditions when it is theoretically sound to use FracMinHash and estimate cosine, and suggested a minimum scale factor that is safe to use. We also presented a fast FracMinHash sketch generator tool `frac-kmc` and benchmarked its running time against `Simka` and `Mash`. We found that when a huge number of small samples are compared, using `frac-kmc` is nearly as fast as `Mash` in wall-clock time. When a number of larger samples are compared, using `frac-kmc` requires more time, although the results produced by `frac-kmc` are more accurate and precise. Our analyses show that when very large sequence files need to be sketched using FracMinHash, using `frac-kmc` can be especially useful.

### 6.2 Further improvements

From a theoretical point of view, the proof technique we used in our theoretical analyses may be applied to other distance/similarity metrics when proving the unbiased expectation proves to be mathematically intractable, and the quantity of interest involves ratios of L2-norms of vectors (such as Bray-Curtis, Chord, Whittaker etc.) From an implementation perspective: the programs we used may be improved and extended in several ways: the code we used to read in MinHash and FracMinHash sketches (generated by `Mash` and `frac-kmc`) is written completely in Python, with not a particular focus on optimization. A well-written C++ implementation may improve things further. The implementation of MurMurHash64 in `frac-kmc` makes use of C++ optimizations, although we did not explore if they can be improved further. `frac-kmc` currently does not support protein *k*-mers (which sourmash does). And finally, instead of using an exact *k*-mer-counter, other approximation-based inexact *k*-mer-counter program may be explored.

## 7 Acknowledgements

We want to thank Marek Kokot and Sebastian Deorowicz for providing us with directions to navigate through the source code of KMC. The work was supported by grant R01GM146462.

